# The UCR Minicore: a valuable resource for cowpea research and breeding

**DOI:** 10.1101/2021.02.09.430326

**Authors:** María Muñoz-Amatriaín, Sassoum Lo, Ira A. Herniter, Ousmane Boukar, Christian Fatokun, Márcia Carvalho, Isaura Castro, Yi-Ning Guo, Bao-Lam Huynh, Philip A. Roberts, Valdemar Carnide, Timothy J. Close

**Affiliations:** Department of Botany and Plant Sciences, University of California Riverside, Riverside, CA 92521, USA; Department of Soil and Crop Sciences, Colorado State University, Fort Collins, CO 80523, USA; Plant Sciences Department, University of California Davis, Davis, CA 95616, USA; Department of Plant Biology, Rutgers University, New Brunswick, NJ 08901, USA; International Institute of Tropical Agriculture, Ibadan, Nigeria; Centre for Research and Technology of Agro-Environmental and Biological Sciences (CITAB), University of Trás-os-Montes and Alto Douro (UTAD), 5000-801, Vila Real, Portugal; Department of Nematology, University of California Riverside, Riverside, CA 92521, USA

## Abstract

Incorporation of new sources of genetic diversity into plant breeding programs is crucial for continuing to improve yield and quality, as well as tolerance to abiotic and biotic stresses. A minicore (the “UCR Minicore”) composed of 368 worldwide accessions of cultivated cowpea has been assembled, having been derived from the University of California, Riverside cowpea collection. High-density genotyping with 51,128 SNPs followed by principal component and genetic assignment analyses identified six subpopulations in the UCR Minicore, mainly differentiated by cultivar group and geographic origin. All six subpopulations were present to some extent in West African material, suggesting that West Africa is a center of diversity for cultivated cowpea. Additionally, population structure analyses supported two routes of introduction of cowpea into the U.S.: (1) from Spain to the southwest U.S. through Northern Mexico, and (2) from Africa to the southeast U.S. via the Caribbean. Genome-wide association studies (GWAS) of important agronomic traits including flowering time, resulted in the identification of significant SNPs for all traits and environments. The mapping resolution achieved by high-density genotyping of this diverse minicore collection allowed the identification of strong candidate genes, including orthologs of the Arabidopsis *FLOWERING LOCUS T.* In summary, this diverse, yet compact cowpea collection constitutes a suitable resource to identify loci controlling complex traits, consequently providing markers to assist with breeding to improve this crop of high relevance to global food and nutritional security.

## INTRODUCTION

Cowpea (*Vigna unguiculata* L. Walp) is a diploid (2*n* = 22) warm-season legume of major importance for food and nutritional security. It provides a major source of dietary protein, fiber, minerals, and vitamins for millions of people in sub-Saharan Africa (SSA), and fodder for livestock. Most of the production is in SSA by smallholder farmers, most of whom are women. It is also grown in many other parts of the world including Latin America, Southeast Asia, the Mediterranean Basin, and the United States (FAOSTAT, www.fao.org). Cowpea is well-known for its adaptation to heat and drought, and to soils with low fertility, making it a successful crop in arid and semi-arid regions where most other crops do not perform as well [1]. However, breeding for increased heat and drought tolerance as well as for key agronomic traits and pest and disease-resistance is crucial as climate changes associated with global warming increase, and given that cowpea is primarily grown in regions that are quite vulnerable to climate change [2,3].

Plant genetic resources constitute the raw material for crop improvement. Cowpea is a genetically diverse crop species divided into five cultivar groups: *unguiculata, biflora, sesquipedalis, textilis* and *melanophthalmus* [4,5]. The two most important cultivar groups are *unguiculata* (grain-type cowpea) and *sesquipedalis* (vegetable-type cowpea, also known as asparagus bean or yardlong bean) [4]. The long- and succulent-podded vegetable cowpea is mostly grown in southeastern Asia while the short-podded grain cowpea is prevalent in Africa and elsewhere [1,6]. In addition to being grown for grain, cowpea is a source of nutritious fodder for livestock in dry savanna regions of sub-Saharan Africa [7]. One important aspect of cowpea agronomy is the time to flowering, as early-maturing types can in many cases be deployed as a strategy to capitalize on shortened periods of optimal growth, thus avoiding late-season drought with its accompanying array of biotic stressors [1].

Diverse cowpea germplasm is available from genebanks around the world, as partially summarized in Genesys (gene.sys-pgr.org/c/cowpea). The largest germplasm collection, comprised of 15,933 accessions, is located at the International Institute of Tropical Agriculture (IITA) in Ibadan, Nigeria. Other large collections, considerably overlapping in content, are at the United States Department of Agriculture – Agricultural Research Service (USDA-ARS) Plant Genetic Resources Conservation Unit (Griffin, Georgia, USA) with 8,242 accessions, the National Bureau of Plant Genetic Resources (NBPGR, New Delhi, India) with 3,704 accessions, and the University of California, Riverside (UCR, California, USA) with about 5,000 accessions. Managing and evaluating large germplasm collections is laborious and costly, and developing core and minicore subsets to facilitate access to the diversity contained in the entire set of accessions is a common practice in most *ex situ* collections [8,9].

Genetic and phenotypic evaluation of diverse collections are needed to fully utilize their potential in breeding programs. Several previous studies have reported on the genetic diversity of cultivated cowpea germplasm. Huynh et al. [10] genotyped 442 cowpea landraces, predominantly from Africa, with a 1,536-SNP GoldenGate assay [11] and identified two major genepools, nominally West versus Southeast Africa. Later, 768 accessions mostly from the USDA cowpea collection were characterized using 5,828 GBS (genotyping-by-sequencing) SNPs, revealing the presence of three major subpopulations [12], again one from West Africa, with the other two subpopulations separating Asian, European and some US accessions from those originating in India, Oceania, other parts of Africa, and the Americas. More recently, the majority of an IITA minicore collection (298 accessions) was genotyped using GBS with 2,276 SNPs, also identifying three major subpopulations [13], but with more dispersion of West and Central African accessions across the three sub-populations. Carvalho et al. [14] genotyped a smaller set of 96 worldwide cowpea accessions emphasizing Iberian Peninsula germplasm, but at a much higher SNP density using the Illumina Cowpea iSelect Consortium Array containing 51,128 SNPs [15]. In this latter work, logical relationships were noted between the genetic composition of accessions and colonial-era movement of germplasm from the Iberian Peninsula to the Caribbean, and from Africa to South America. In general, it is evident that there is much yet to be clarified regarding the spread of cowpeas worldwide, and the extent to which certain genetic variants have taken hold in different regions at different times.

Here, we report the development of a minicore (henceforth referred as the “UCR Minicore”) composed of 368 domesticated cowpeas selected from a larger set of ~5,000 accessions comprising the UC Riverside cowpea collection. This minicore was genotyped using the 51,128-SNPs Cowpea iSelect Consortium Array [15] and the genotypic information was used to gain a better understanding of the genetic diversity of domesticated cowpea. Although this is the first summary report and full SNP data release for the UCR Minicore, this material has been utilized for more focused work on seed coat color [16], seed coat pattern [17], seed size [18], bruchid resistance [19], plant herbivore resistance [20] and pod shattering [21]. This study evaluated additional traits of agronomic importance including flowering time, dry pod weight, dry fodder weight and pod load score. Genome-wide association studies (GWAS) were conducted, identifying significant SNPs and candidate genes associated with each of these traits.

## MATERIALS AND METHODS

### Plant materials and phenotyping

A total of 668 accessions representing a core subset from the University of California Riverside (UCR) cowpea germplasm collection were genotyped with an Illumina GoldenGate assay [11] (1,536-SNPs) in previous studies [10,22]. This SNP information, coupled with available phenotypic and passport data, was used to choose a smaller subset of non-redundant accessions, each of which was highly homozygous, collectively representing the genetic and phenotypic diversity of the larger core collection. A few additional accessions nominated by cowpea researchers and breeders based on regions of origin not represented among the core collection and possessing some specific traits were also included. The minicore of 368 accessions includes landraces and breeding materials from 50 countries in Africa, Asia, North and South America, Europe, and Australia (Table S1) encompassing members of the cultivar groups *unguiculata* and *sesquipedalis.* Individual plants were grown from each of the 368 accessions in a greenhouse at UCR for genotyping (see section *2.2*) and seed production. Subsequent seed increases for distribution and phenotypic evaluations used seeds directly descended from these genotyped plants.

The UCR Minicore was phenotyped for days to flowering (DTF) in California (USA) under long-days at the UCR Citrus Research Center and Agricultural Experiment Station in Riverside (CA) during the summers of 2016 and 2017, as well as under short days at the UCR Coachella Valley Agricultural Research Station in Thermal (CA) during the autumn of 2016. For the Riverside summer planting, daylight hours (the time between sunrise and sunset) ranged from 14.4 h on 21 June to 11.9 h on 30 September. For the Thermal autumn planting, daylight hours ranged from 12.8 h on 1 September to 9.9 h on 21 December. Due to limited seed availability, 50 seeds of each accession were planted in single rows of 5.5 – 6.1 m long with 0.75 m spacing between rows at both locations. Scoring of the minicore was also conducted at two locations in Nigeria during 2017. The minicore was sown in August 2017 at the International Institute of Tropical Agriculture (IITA) experimental field of Malamadori and Minjibir, near Kano, Nigeria. An alpha lattice design with three replications was used. Each accession was assigned to a plot of 2 m length. The distance between two consecutive plots was 0.75 m while two plants within a row were separated by 0.20 m. The fertilizer NPK (15-15-15) was applied two weeks after planting at the rate of 100 kg/ha. To control insect pests, the trial was sprayed with the insecticide Kartodim (Dimethoate 300g + Lambda-cyhalothrin 15g) at the rate of 1.2 l/ha four times: once at each of vegetative and at flower opening and twice during pod maturing. Manual weeding was performed twice to control weeds. At all locations, DTF was scored as the number of days from planting to when 50% of plants had at least one flower opened. Pod load score was recorded at plant maturity using a 1-3 scale with 1 for high pod load (80-100% of peduncles had 2-3 pods per peduncle), 2 for moderate pod load (80-100% of peduncles had 1-2 pods per peduncle) and 3 for poor load score (80-100% of peduncles had 1 or less pod per peduncle). At maturity, dry pods were harvested, and the other above-ground parts of the plant (leaves, twigs and stems) were cut and rolled up. Both the pods and the fodder were sun dried for one week to determine dry pod weight and fodder weight respectively.

### SNP genotyping and data curation

Genomic DNA was extracted from young leaves of individual plants using DNeasy Plant Kit (Qiagen, Valencia, California, USA). The Cowpea iSelect Consortium Array including 51,128 SNPs [15] was used to genotype each DNA sample at the University of Southern California Molecular Genomics Core (Los Angeles, California, USA). SNPs were called in GenomeStudio (Illumina Inc., San Diego, California, USA) and manually curated to remove those with high levels (> 25%) of missing data and/or heterozygous calls. The highest percentage of heterozygosity for any accession was 3.83% (Cp 4906), and 96% of the accessions had heterozygosity levels below 1% (Table S2). The final dataset included 48,425 polymorphic SNPs, 47,334 with known physical positions [23], on the 368 minicore samples (Table S2).

### Genetic diversity, population structure and linkage disequilibrium analyses

Expected heterozygosity (*He*) and nucleotide diversity (π) values were calculated for all 48,425 SNPs in the minicore as a measure of genetic diversity. *He* was calculated as 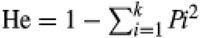, where *P_i_* is the frequency for the *i*^th^ allele among a total of *k* alleles. π was evaluated as in Xu et al. [6].

The admixture model implemented in STRUCTURE v2.3.4 [24] was used to infer population structure of the UCR Minicore. Only SNPs with minor allele frequency (MAF) ≥ 0.05 were included in the analysis. STRUCTURE was first run three times for each hypothetical number of subpopulations (*K*) between 1 and 10, with a burn-in period of 10,000 followed by 10,000 Monte Carlo Markov Chain (MCMC) iterations. LnP(D) values were plotted and Δ*K* values were calculated according to Evanno et al. [25] to estimate the optimum number of subpopulations. Plots were generated with Structure Harvester [26] (Figure S1). Then, a new run using a burn-in period of 100,000 and 100,000 MCMC iterations was conducted for the estimated *K* to assign accessions to subpopulations based on a membership probability ≥ 0.80. In addition, principal component analysis (PCA) and linkage disequilibrium (LD) analysis were conducted on the same SNP set using TASSEL v5 [27].

Linkage disequilibrium (LD) was estimated for each chromosome as the correlation coefficient *r*^2^ between pairs of SNPs, using SNPs with MAF ≥ 0.05 (42,711 SNPs). The decay of LD over physical distance was investigated by plotting pair-wise *r^2^* values and generating a locally weighted scatterplot smoothing (LOESS) curve in R. LD decay distance was determined when *r*^2^ fell to the critical threshold estimated from the 95^th^ percentile *r*^2^ distribution for unlinked markers (*r*^2^ = 0.15).

### Genome-wide association studies (GWAS)

The mixed linear model [28] implemented in TASSEL v.5 [27] (http://www.maizegenetics.net/tassel) was used for GWAS, with a population structure matrix (for *K* = 6) and a kinship coefficient matrix accounting for population structure and the relatedness of accessions, respectively. A false discovery rate (FDR; [29]) threshold of 0.01 was used to identify significant associations. The percentage contribution of each SNP to the total phenotypic variation was calculated using marker *R*^2^ values from TASSEL multiplied by 100. Candidate genes within significant regions were identified from annotations of the cowpea reference genome v1.1 [23] (https://phytozome.jgi.doe.gov/).

## RESULTS

### Diversity and linkage disequilibrium in the UCR Minicore

The UCR Minicore was developed to represent available genetic and phenotypic diversity of cultivated cowpea while maintaining a sample size that can be managed by most researchers and breeders for evaluating traits of interest. This minicore is composed of 368 accessions from 50 countries including 242 landraces, 98 breeding lines, three accessions categorized as “weedy”, and 25 accessions that are uncategorized (Table S1). It largely overlaps with the IITA minicore, with 233 accessions in common (Table S1), at least by name though not necessarily by genetic identity. The UCR Minicore showed great phenotypic diversity in pod and seed types and colors, flowering time and maturity, leaf shape, and plant architecture, among other traits. Figure 1 illustrates some of the morphological variation existing in the UCR Minicore.

**Figure 1.**
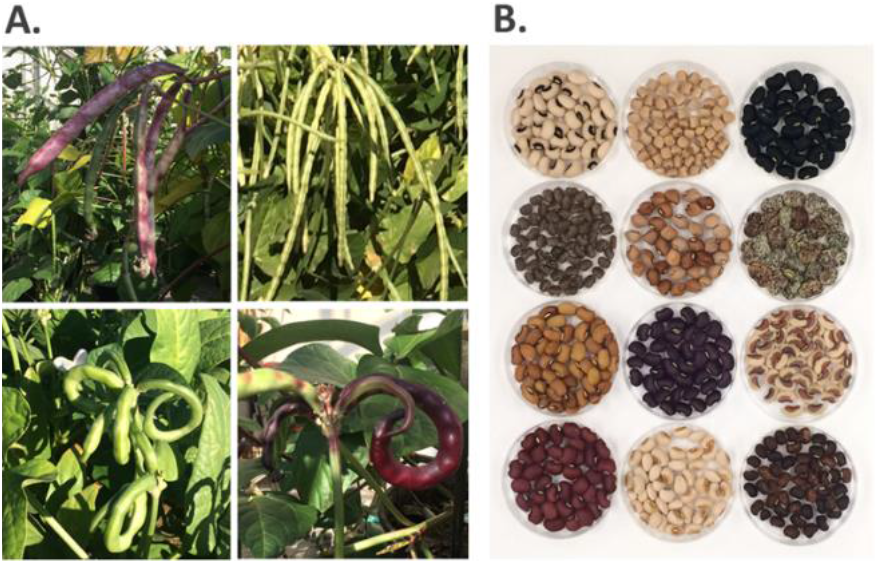
Phenotypic diversity in the UCR Minicore. Examples of diversity in (A) pod color and morphology and (B) seed coat color and pattern.

Genotyping with the 51,128-SNP array [15] enabled analyses of genetic diversity in the minicore. 94.7% of those SNPs (48,425) were polymorphic in the dataset. Expected heterozygosity (*He*) and nucleotide diversity (π) were calculated and both averaged 0.313 (the maximum for a biallelic SNP is 0.5).

The LD decay distance was also investigated for each cowpea chromosome using SNPs with MAF ≥ 0.05. The decay of LD with physical distance differed among chromosomes and ranged from 809 kb on Vu05 to 4,705 kb on Vu10 (Figure S2). In addition to Vu05, the LD decay distance was shortest in Vu08 (813 kb) and Vu07 (1,048 kb), while Vu01 and Vu04 had the largest decay distances after Vu10 (3,803 kb and 3,286 kb, respectively; Figure S2). On average there was one SNP per 14.9 kb, indicating sufficient marker density for GWAS.

### Population structure of the UCR Minicore

A total of 42,711 SNPs with MAF ≥ 0.05 were used to evaluate population structure in the UCR Minicore. STRUCTURE [24] was run for *K* = 1–10 and the inspection of both the estimated log probabilities of the data and the *ΔK* values calculated as in Evanno et al. [25] supported the presence of six genetic subpopulations (Figure S1). Accessions were assigned to each subpopulation using a membership coefficient ≥ 0.8 (accessions with membership coefficients less than 0.8 were considered “admixed”; Table S1). PCA also showed a clear separation of the six subpopulations on the first three principal components (Figure S3).

Subpopulation 1 included 31 accessions mostly from West Africa, 29 of which are landraces (Table S1). Subpopulation 2 was the smallest subpopulation by number of accessions: it included 12 breeding lines developed at the International Institute of Tropical Agriculture (IITA) in Nigeria. Subpopulation 3 was composed mostly of landraces from Mediterranean countries including Egypt, Italy, and Portugal as well as accessions from California and Puerto Rico (Table S1). Subpopulation 4 included 27 accessions that are mostly landraces from India, China and Papua New Guinea, among other countries. Passport data and visual examination of accessions in Subpopulation 4 indicated that they belong to the *sesquipedalis* cultivar group (yardlong beans). Subpopulation 5 contained 32 accessions from West Africa, most of which are landraces (Table S1). The largest subpopulation was Subpopulation 6 (69 accessions), which was composed primarily of landraces from Southeastern Africa. The remainder of the accessions (165) were considered “admixed” (Table S1).

Figure 2 shows the worldwide distribution of the six subpopulations, with each pie plot representing the proportion of the six subpopulations contributing to accessions in each country. Accessions within the USA were divided further based on the state they belonged to. Each accession from California is represented on the map, while the rest of the USA accessions were grouped together due to the lack of passport information about the state of origin for most of them. Their cultivar names, however, traced to U.S. southern states (Table S1). The figure graphically shows that while all subpopulations are present in West Africa, three of them (1, 2 and 5) are predominant. Accessions from Southeastern Africa have ancestry primarily from Subpopulation 6, while most germplasm from Mediterranean countries belonged to Subpopulation 3. Interestingly, a closer look at the germplasm from the USA indicates that accessions from California have ancestry mostly from Subpopulation 3, while accessions from other U.S. states are predominantly Subpopulation 6 (Figure 2). Subpopulation 4 is primarily from countries in Asia and Oceania.

**Figure 2.**
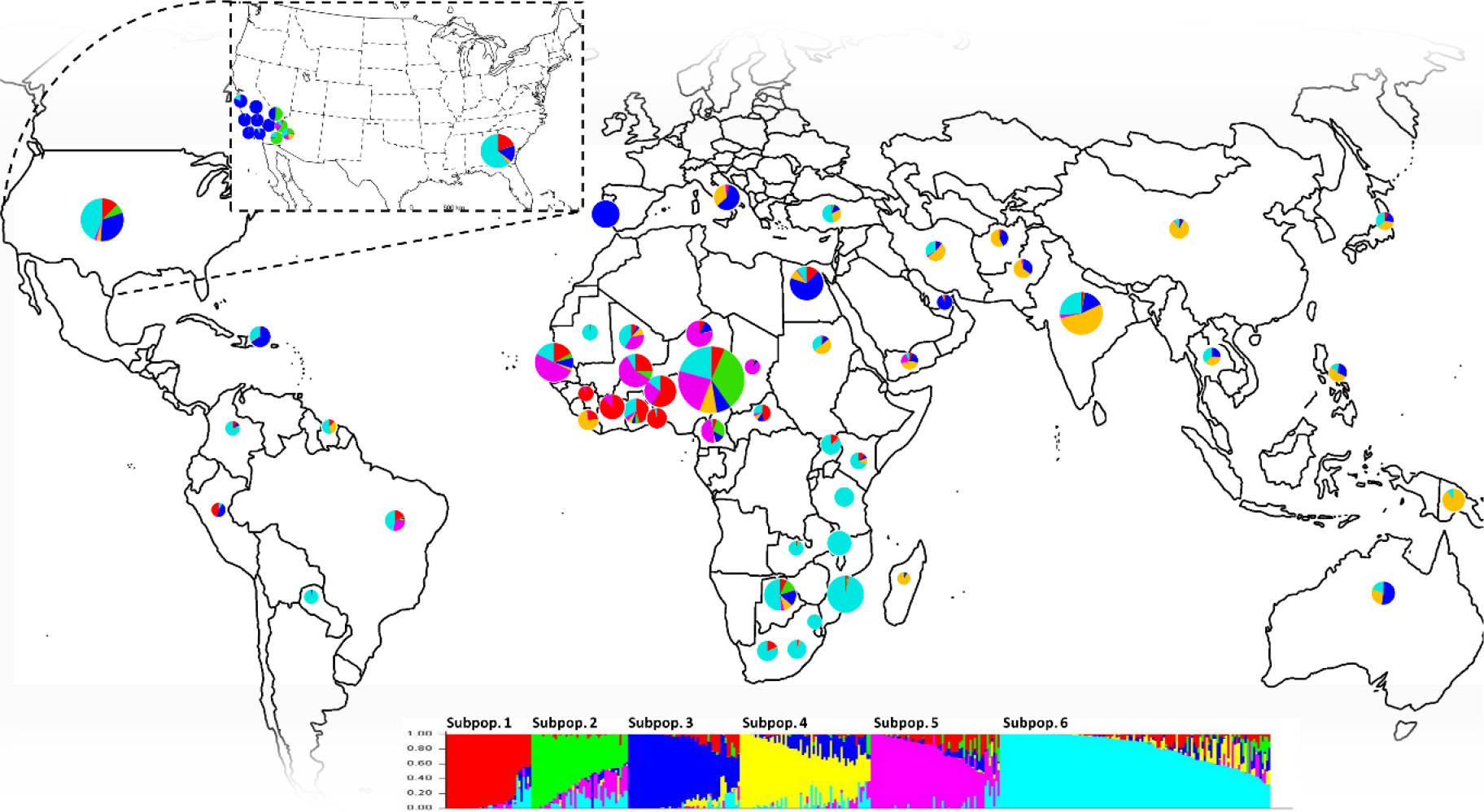
Population structure in the UCR Minicore. The geographical distribution of the accessions belonging to the minicore is shown, with pie charts representing the average proportions from each genetic subpopulation for samples in each country. Pie chart sizes are proportional to the number of samples in each country and range from 1 to 80 accessions. Accessions within the U.S. were further divided as “California” and “other states/unknown”, as most accessions in this second group did not have state information but their cultivar names traced to U.S. southern states. The plot of ancestry estimates for *K*=6 is shown at the bottom, with each bar representing the estimated membership coefficients for each accession in each of the six subpopulations (represented by different colors).

### GWAS of agronomic traits

GWAS was conducted for four different agronomic traits including days to flowering (DTF), pod load score, dry pod weight, and dry fodder weight using 42,711 SNPs with MAF ≥ 0.05 (see Materials and Methods). DTF was evaluated in five different environments in the USA and Nigeria, three of which are considered short-day environments (< 12h of daylight) while the other two are considered long-day environments (> 12h of daylight) (Table 1, Materials and Methods). Pod load score, dry pod weight, and dry fodder weight were evaluated in one environment in Nigeria. Significant marker-trait associations were identified for all traits and environments (Figure 3 and 4; Table 1 and 2; Table S3 and S4).

**Figure 3.**
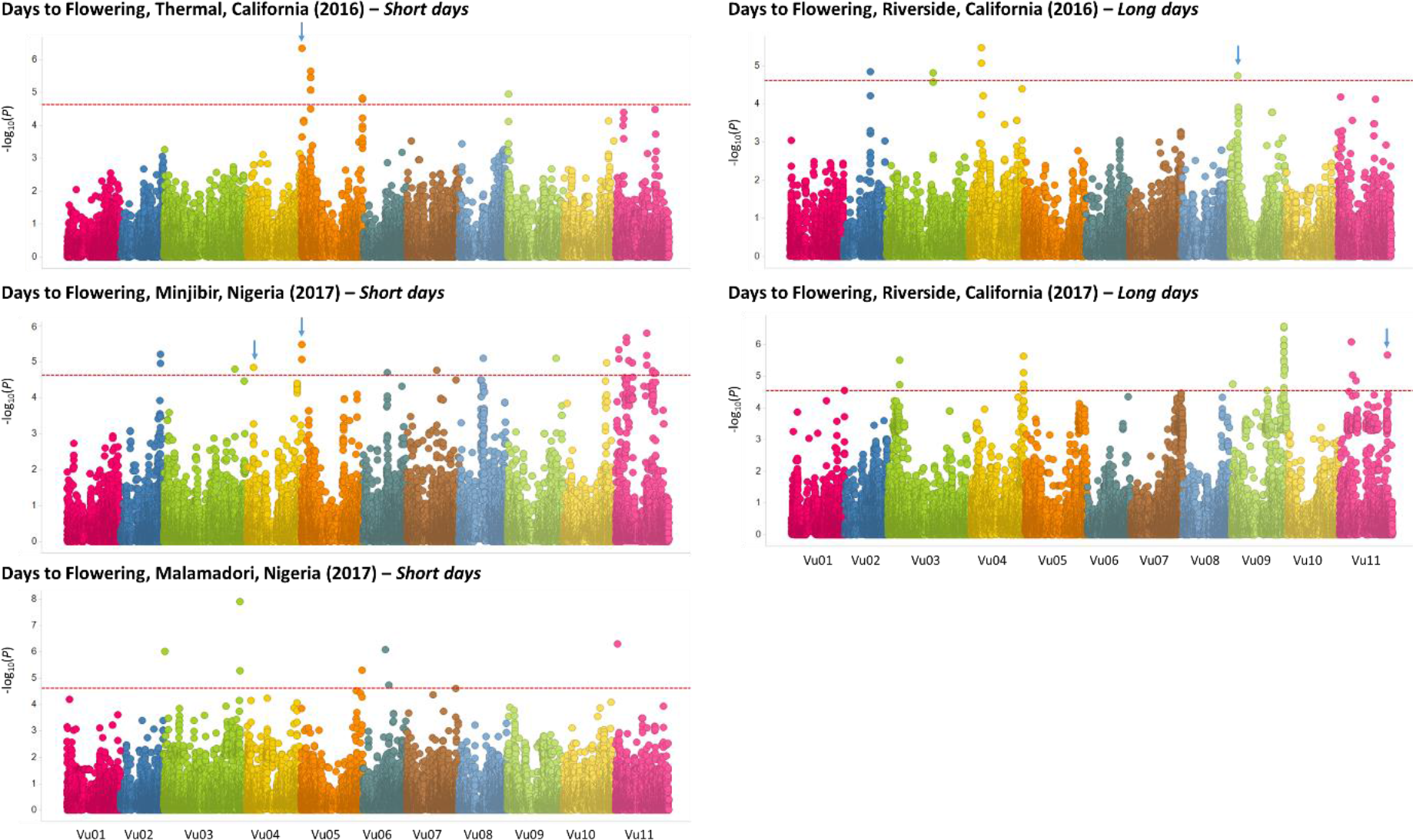
Manhattan plots of GWAS on days to flowering in five different environments at four locations. −log10 (p-values) are plotted against physical positions on the cowpea reference genome v1.1 [23]. The dashed red line in each plot indicates the 0.01 FDR-corrected threshold, ranging from 4.54 for Riverside, California (2017) to 4.62 for both Thermal, California (2016) and Minjibir, Nigeria (2017). Blue arrows represent known QTLs for DTF.

**Figure 4.**
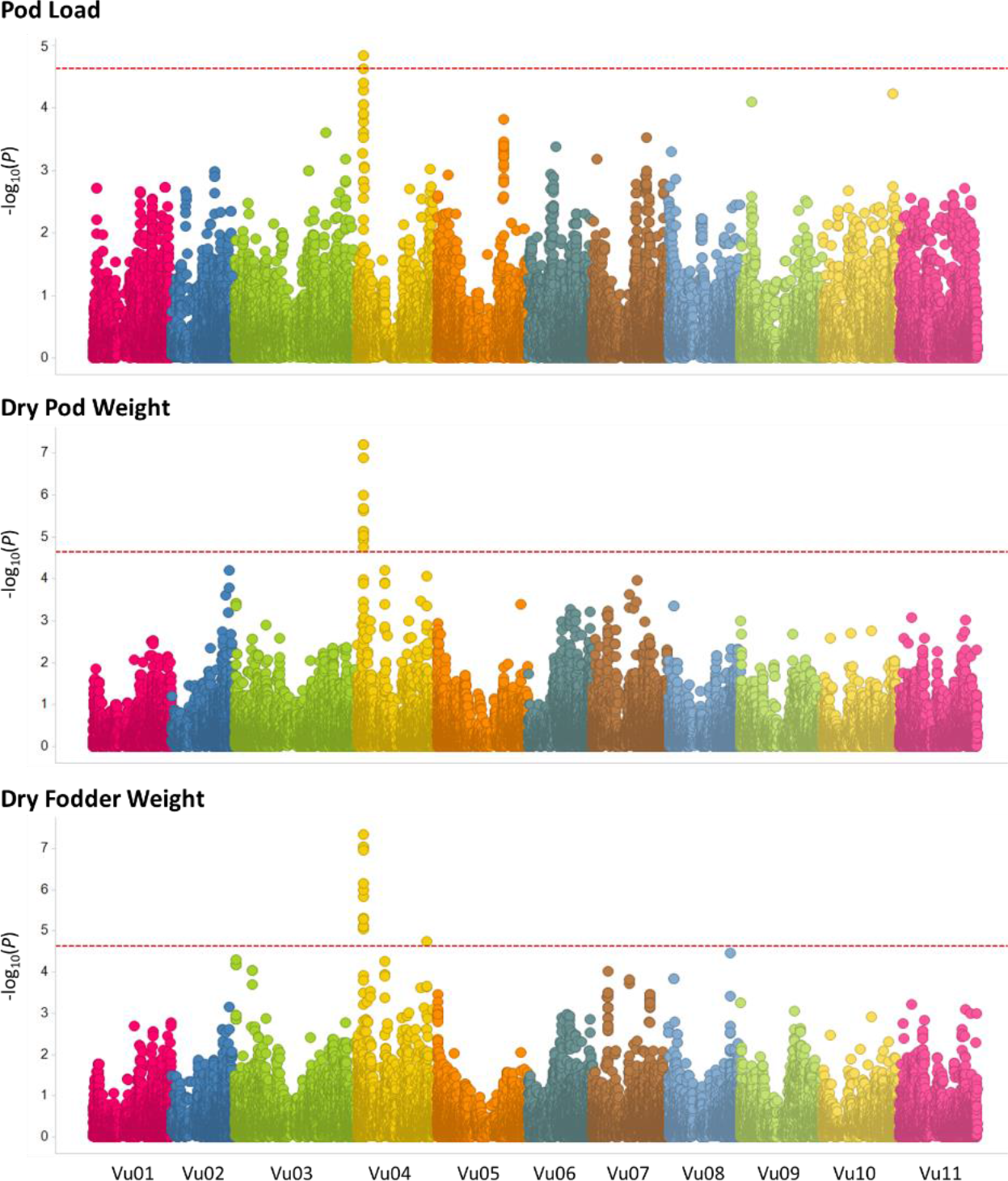
Manhattan plots of GWAS on pod load score, dry pod weight, and dry fodder weight. −log10 (p-values) are plotted against physical positions on the cowpea reference genome [23]. The dashed red line in each plot indicates the 0.01 FDR-corrected threshold (4.62 for dry fodder weight and 4.63 for both pod load score and dry pod weight).

**Table 1.**
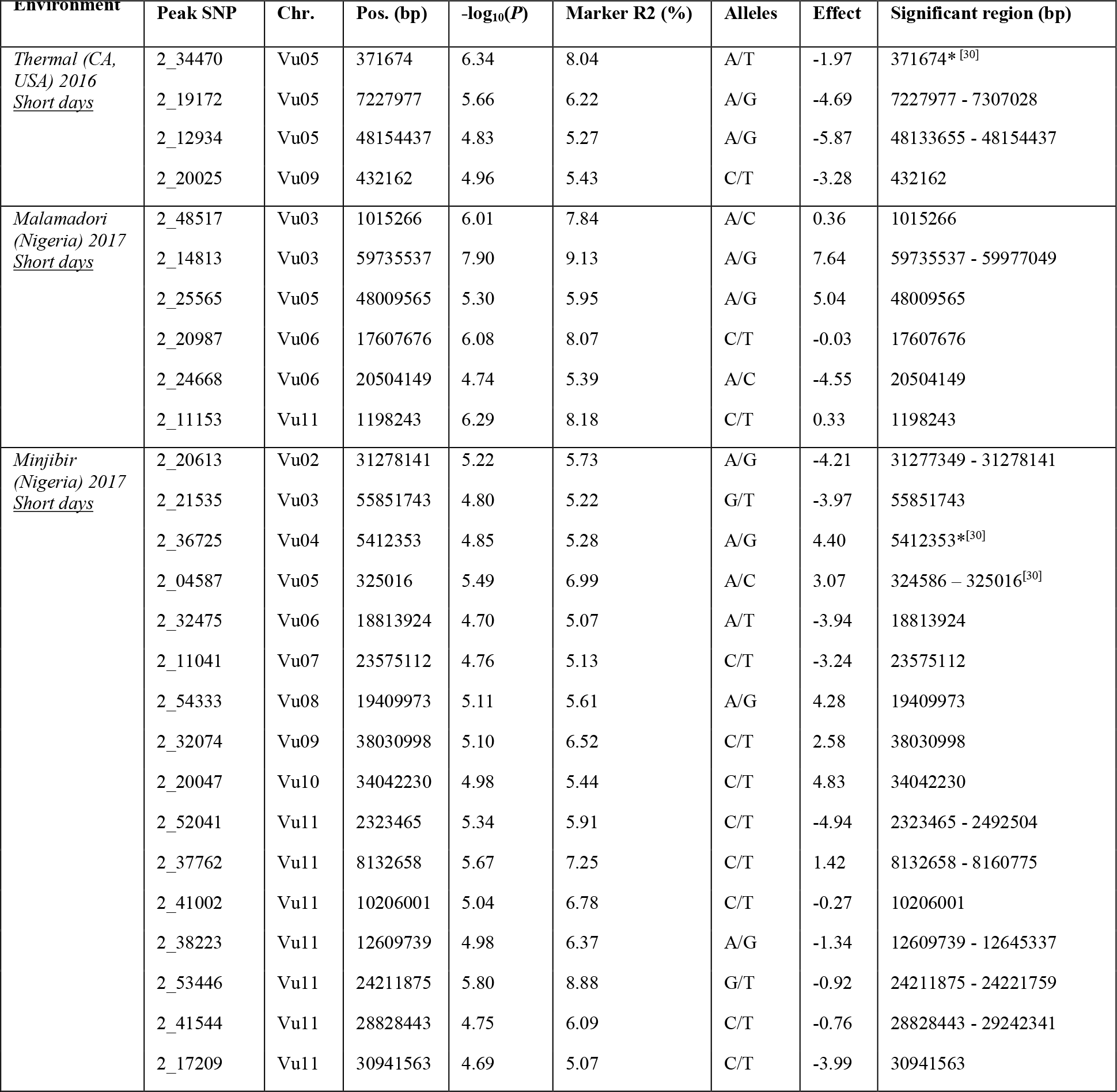
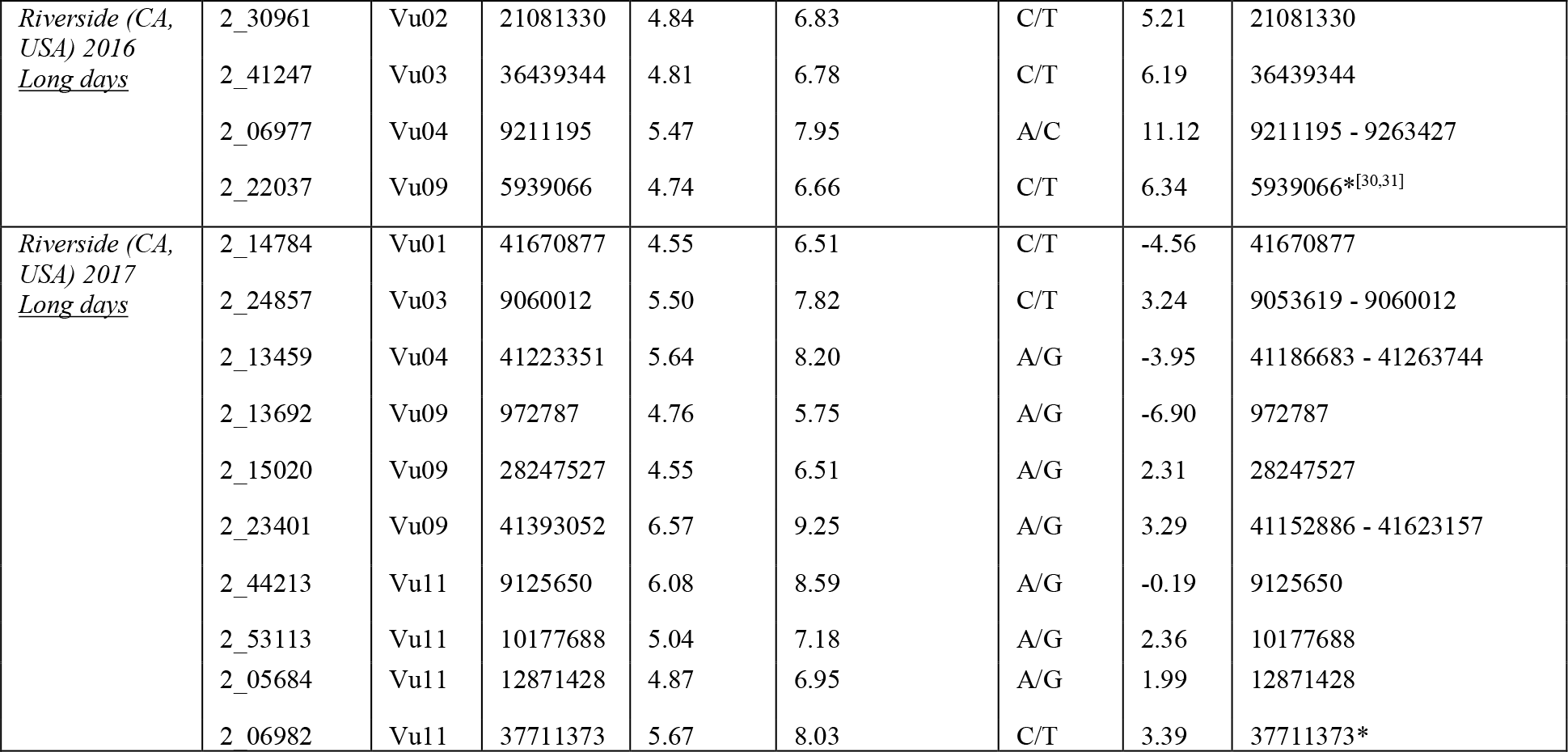
Peak SNPs associated with DTF under five different environments. Asterisk on significant region (bp) indicates previously known QTL for DTF, as discussed in the text.

**Table 2.** Peak SNPs associated with dry fodder weight, dry pod weight and pod load.

| Trait | Peak SNP | Chr. | Position (bp) | $-\log_{10}(P)$ | Marker R<sup>2</sup> (%) | Alleles | Effect | Significant region (bp) |
| --- | --- | --- | --- | --- | --- | --- | --- | --- |
| *Pod Load Score* | 2_05691 | Vu04 | 3403047 | 4.84 | 6.32 | G/T | -0.15 | 3335635 - 3403521 |
| *Dry Pod Weight* | 2_06769 | Vu04 | 3278892 | 7.21 | 8.85 | C/T | 29.60 | 3278476 - 3403521 |
| *Dry Fodder Weight* | 2_05693 | Vu04 | 3403521 | 7.34 | 10.39 | C/T | 20.87 | 3278476 - 3403521 |
|  | 2_02590 | Vu04 | 37245973 | 4.75 | 6.64 | A/G | -9.43 | 37245973 |

### Days to flowering

Considering all five environments, 91 significant SNPs at 40 genomic regions were identified for DTF (Figure 3, Table 1, Table S3). Among those 40 significant QTLs, 26 were associated with DTF under short days in Thermal (California, USA) and both Malamadori and Minjibir (Nigeria), while 14 were associated with DTF under long days in Riverside (California, USA) (Figure 3, Table 1, Table S3). The phenotypic variation of significant SNPs ranged from 5 - 9% (Table 1, Table S3).

Five of the significant genomic regions corresponded to previously reported QTLs for DTF (Figure 3; Table 1; [30,31]) while the rest are novel (Table 1; Figure 3). Those five QTLs were investigated further by considering gene model annotations in Phytozome (phytozome.jgi.doe.gov/) and the Legume Information System (LIS; www.legumeinfo.com). The resolution of QTL positions was improved in the present work using higher-density genotyping than previously, together with a larger and more diverse set of accessions.

Several candidate genes stood out for these five QTLs, as follows. One of the previously reported QTL was identified in Thermal 2016 and is located at the beginning of Vu05 (371,674 bp; Table 1). Just 47 kb away from that position, another QTL was identified in Minjibir 2017 between 324,586 bp and 325,016 bp (Table 1). These coordinates coincide with a QTL for DTF identified in a cowpea MAGIC population also under short days [30] and hence they are likely to represent the same QTL. Only six genes were found within this region (324,586 - 371,674 bp) including a cluster of four genes (*Vigun05g004000, Vigun05g004100, Vigun05g004200* and *Vigun05g004300*) encoding the flowering locus protein T. Another of these five DTF QTLs was identified by a SNP at 5,412,353 bp on Vu04 under short days in Minjibir in 2017 (Table 1; Figure 3). This position is contained within a QTL for flowering time under short days identified in the cowpea MAGIC population by Huynh et al. [30]. Genes near the significant SNP (2_36725) include *Vigun04g057300,* encoding a circadian clock coupling factor ZGT, ~200 kb from this SNP. *Vigun04g057300* is an ortholog of the Arabidopsis *Empfindlicher im Dunkelroten Licht 1* (*EID1*) gene involved in the regulation of phytochrome-A light signaling [32].

Another of the five previously reported QTLs affects flowering time under long-days. A single significant SNP (2_22037) was identified at 5,939,066 bp on Vu09 in Riverside in 2016 (Table 1; Figure 3). This position matches a QTL for DTF identified previously in the same environment in the MAGIC population [30] as well as a flowering time QTL identified in a RIL population derived from a cultivated x wild cross (*CFt9;* [31]). *Vigun09g059700,* which encodes a MADS-box protein and is orthologous to the Arabidopsis *AGAMOUS-LIKE 8* (*AGL8*) gene also known as *FRUITFULL (FUL),* was identified 140 kb from this SNP. Lastly, another of these five QTLs was associated with flowering under long days. This locus was detected at position 37,711,373 on Vu11 for the Riverside 2017 environment (Table 1; Figure 3), which is contained within a flowering time QTL previously identified in the same environment using the cowpea MAGIC population [30]. Two cowpea genes, *Vigun11g169600* and *Vigun11g169400,* encoding AP2/B3-like transcription factor family proteins were identified at 128 kb and 139 kb from the peak SNP, respectively. These genes are orthologous to the Arabidopsis *REDUCED VERNALIZATION RESPONSE 1 (VRN1),* a major gene involved in regulation of flowering and vernalization response.

In addition to these previously reported cowpea flowering time QTLs, it is worth mentioning three other QTLs that contain or are located near genes with clear roles in flowering. One is the QTL identified in Thermal in 2016 between 7,227,977 - 7,307,028 bp on Vu05 (Table 1; Figure 3). The cowpea gene *Vigun05g077400,* encoding a MADS-box protein orthologous to the Arabidopsis AGAMOUS-like 20 (AGL20) gene, is located 43.7 kb from the peak SNP (2_19172) for this QTL. The second one is a QTL region spanning 470 kb (41152886 to 41623157 bp) on Vu09 that was identified in Riverside in 2017, a long-day environment. This region contains 69 genes, among which *Vigun09g244300*, encoding a protein of the BES1/BZR1 family, was found. This gene is orthologous to the Arabidopsis *BRASSINAZOLE-RESISTANT 1 (BZR1),* which positively regulates the brassinosteroid signaling pathway [33]. The third QTL of interest was also identified in Riverside in 2017; it was located on Vu03 between 9,053,619 and 9,060,012 bp (Table 1; Figure 3). The cowpea gene *Vigun03g104200,* which is an ortholog of the soybean *E6*, a main flowering and maturity gene, was identified 21 kb upstream of the peak SNP (2_24857).

### Pod load score, dry pod weight, and dry fodder weight

Pod load score, dry pod weight and dry fodder weight were evaluated in Minjibir in 2017 (Nigeria). Correlations between these traits were calculated using Pearson’s correlation coefficient. A strong positive correlation was observed between dry pod weight and dry fodder weight (0.93), while a moderate negative correlation was observed between pod load score and dry fodder weight (−0.45) as well as between pod load score and dry pod weight (−0.48). Note that lower pod load score numbers were given to plants with higher pod loads (see Methods).

A single QTL was identified for pod load score and dry pod weight on Vu04, while two QTLs on Vu04 were identified for dry fodder weight (Table 2; Figure 4; Table S4). Interestingly, the major QTL for dry fodder weight coincides with the QTLs for pod load score and dry pod weight (Table 2; Figure 4; Table S4). The colocation of these QTLs together with the correlations between the three traits suggests pleiotropic effects of a single gene or the existence of closely linked genes.

Genes within this common QTL region which spans 125 kb were explored. Eleven genes were identified, seven of which encode members of the cyclic nucleotide-gated ion channel (CNGC) family of proteins. Four of those seven genes (*Vigun04g039300*, *Vigun04g039400*, *Vigun04g039800* and *Vigun04g039900*) are orthologs to the Arabidopsis *CNGC20* gene, also called *CNBT1* (cyclic nucleotide-binding transporter 1), while the other three (*Vigun04g039500*, *Vigun04g039600* and *Vigun04g039700*) are orthologs to the Arabidopsis *CNGC19* gene. The encoded ion channel proteins mediate signaling pathways involved in responses to abiotic and biotic stresses [34], including response to herbivores, nematodes and heavy metals among others [35–37] (see Discussion).

## DISCUSSION

### Population structure and historical crop dispersal

The UCR Minicore and its associated high-density SNP data (51,128 SNPs) constitute a powerful combination of material and information resources to support genetic diversity analyses of cultivated cowpea. Six subpopulations were identified in this minicore, all of which were represented to at least some extent in West African material (Figure 2). This indicates that West Africa is a center of diversity for cultivated cowpea, as suggested by previous studies [13,38,39]. Five of the six subpopulations were composed mostly of landraces, while Subpopulation 2 included cowpea lines only from IITA’s cowpea breeding program. Based on how the six subpopulations split at different *K* numbers, as illustrated in Herniter et al. [40], it appears that Subpopulation 2 is the result of crosses between materials from Subpopulations 1 and 5. Subpopulations 1 and 5 are composed almost exclusively of West African accessions. A closer inspection of landraces from Subpopulations 1 and 5 revealed consistent differences in several traits. For example, Subpopulation 1 accessions flowered earlier than those in Subpopulation 5 (8 days earlier on average in short-day environments) and had decreased photoperiod sensitivity (most Subpopulation 5 accessions did not flower under long-days) (Table S1). Other work reported that Subpopulation 1 had much more pod shattering than Subpopulation 5 accessions [21]. In addition, all accessions in Subpopulation 1 have smooth seed coats, while most in Subpopulation 5 have rough coats (data not shown). Seed coat texture is an important quality trait that influences seed end-use; rough seed-coated varieties are preferred for food preparations requiring seed coat removal (e.g., making of Akara) as rough seed coats can be easily removed after soaking, while varieties with smooth seed coats are often preferred when cowpea is consumed as boiled intact seed [41]. Also, rough seed coat types imbibe water quicker and generally have reduced cooking times compared to smooth seed coat types.

Genotyping of the UCR Minicore has shed light on the history of cowpea in USA. Landraces and their breeding derivatives from California belong to Subpopulation 3, which is composed mostly of landraces from Mediterranean countries, while accessions from other U.S. states were predominantly from Subpopulation 6, which is composed mostly of landraces from Southeastern Africa. This population structure, together with textual evidence summarized by Herniter et al. [40], is consistent with a global dispersal of cowpea from its centers of domestication in West and East Africa along historical trade routes. For the USA, cowpea appears to have arrived through at least two distinct introduction routes. It is believed that in the US Southwest, cowpea was first introduced by the Spanish explorer Hernando de Alcorón in 1540 going northward from Mexico, possibly followed in the late 1600’s by the Jesuit monk Eusebio Kino, including accessions that were popular in the Mediterranean basin during those times [40,42]. In the Southeastern United States, cowpea seems to have been brought on slave ships, perhaps as provisions. The two distinct introduction routes of apparently genetically distinct cowpeas contrast with older studies predating genotyping, which have often assumed a single introduction. This conjecture of more than one route of cowpea introductions into the USA, and genetic distinctions between them that now are evident, is further supported by the findings of Carvalho et al. [14], which showed relationships between a Cuban and Mediterranean accessions, and between sub-Saharan African and South American accessions, which represent yet another route of colonial-era dispersal of cowpeas.

Although the germplasm utilized in this study largely overlaps with that utilized by Huynh et al. [10], only two genetic clusters were identified in that previous study which correspond to the two major “West Africa” and “Southeastern Africa” gene pools. As shown by Herniter et al. [40] utilizing this same UCR Minicore material, at *K*=2 Subpopulation 6 (Southeastern Africa gene pool) splits from the rest of the subpopulations, confirming that as a primary genetic differentiation between subpopulations of domesticated cowpeas. The smaller number of SNPs that were available for Huynh et al. [10] to conduct population structure analyses, together with an emphasis on African landraces among accessions genotyped, most likely precluded further subdivision of the West African subpopulation.

Carvalho et al. [14] has been the only previous study to genotype a set of cowpea accessions at a high density (51,128 SNPs). Although their focus was on genetic diversity of Iberian Peninsula cowpeas, results from that study largely agree with the findings reported in the present work. In particular, Subpopulations 1, 2, and 3 from the study of Carvalho et al. [14], correspond to Subpopulations 4, 3 and 6 reported here, respectively. Their fourth subpopulation was composed only of four accessions from West Africa. This small representation of germplasm from West Africa certainly would preclude the detection of additional subpopulation structure present in that region.

### Flowering time

Flowering time is one of the most important agronomic traits, which affect environmental adaptation and yield potential. It is controlled by multiple genes via different pathways, and it is influenced by environmental conditions [43].

Cowpea is generally a short-day plant and, although some accessions are day-neutral (photoperiod-insensitive), many cowpea genotypes are photoperiod sensitive and show a delay in flowering under long day conditions [44,45]. The degree of photoperiod sensitivity can vary between accessions and is influenced by temperature [45]. Understanding the underlying genetic factors of cowpea flowering time is important for the use of diverse germplasm to customize varieties for different environments.

GWAS using the UCR Minicore has enabled the identification of many loci and meaningful candidate genes associated with flowering under both short and long days. Five of the significant regions coincide with QTLs reported in previous studies [30,31]. Overall, these results are consistent with polygenic control of flowering time in cowpea.

One of the main flowering time regions identified under short-day conditions (and two different environments) was on Vu05. A cluster of four genes (*Vigun05g004000, Vigun05g004100, Vigun05g004200* and *Vigun05g004300*) annotated as *FLOWERING LOCUS T* (*FT*) are in this region. *Vigun05g004000* and *Vigun05g004100* are orthologous to the Arabidopsis flowering gene *TWIN SISTER OF FT (TSF; AT4G20370.1),* which is a close relative of *FT,* whereas *Vigun05g004200* and *Vigun05g004300* are orthologs of the Arabidopsis *FT* (*AT1G65480.1*) gene. *TSF* and *FT* are main floral pathway integrators and play overlapping roles in the promotion of flowering [43,46,47]. They share a similar mode of regulation, and overexpression of both *FT* and *TSF* results in precocious flowering [46,47]. The identification of multiple copies of *FT* genes in the cowpea reference genome [23] suggests that copy number variation could play an important role in the regulation of flowering time in cowpea, as reported for other crops [48,49].

Another main flowering time locus identified under short days was located near *Vigun04g057300,* which is an ortholog of the Arabidopsis gene *EID1. EID1* encodes an F-box protein that is a negative regulator in phytochrome A (phyA)-specific light signaling [50]. Mutations in *EID1* causing the deceleration of the circadian clock have been selected during tomato domestication [51]. In Arabidopsis, *EID1* mutations caused alterations in flowering induction [32].

A third QTL identified under a short-day environment on Vu05 was near *Vigun05g077400*, which is an ortholog of the Arabidopsis *AGAMOUS-like 20* (*AGL20*) gene. *AGL20* is an integrator of different pathways controlling flowering and is considered a central component for the induction of flowering [52,53]. Overexpression of *AGL20* in Arabidopsis suppresses late flowering and delays phase transitions from the vegetative stages of plant development [52,53].

Under the long-day conditions of Riverside (CA, USA), most of the accessions in the UCR Minicore showed a delay in flowering, and about one fourth of the minicore did not flower (Table S1). Under long days, a major flowering time QTL was identified on Vu09 near *Vigun09g059700*, an ortholog of the Arabidopsis *AGL8/FUL* gene. This gene encodes a MADS-box family transcription factor that regulates inflorescence development and is negatively regulated by *APETALA1* in Arabidopsis [54]. In addition, *AGL8/FUL* promotes floral determination in response to far-red-enriched light [55]. Loss of *AGL8/FUL* function in Arabidopsis caused a delay in flowering time [56,57], suggesting that the cowpea ortholog of *FUL* (*Vigun09g059700*) could similarly be involved in the response to photoperiod.

Another region associated with flowering under long day was identified near genes *Vigun11g169600* and *Vigun11g169400*. Both genes encode AP2/B3-like transcription factors that are orthologs to the Arabidopsis *VRN1* gene. *VRN1*, a close homolog of *FUL*, promotes flowering after prolonged cold (vernalization) in different plant species [58–62]. It is intriguing to identify a vernalization-pathway gene in a warm-season legume that does not need to undergo vernalization before flowering. However, vernalization pathway genes have been identified in other warm-season plants including soybean [63]. In particular, *Glyma11g13220,* which was a homolog of the Arabidopsis *VRN1*, was responsive to photoperiod as well as to low temperatures in soybean. This vernalization pathway gene could also be functional in cowpea.

The cowpea gene *Vigun09g244300,* located on Vu09 and encoding a protein of the BES1/BZR1 family, is another strong candidate gene for days to flowering under long-day conditions. *Vigun09g244300* is an ortholog of the Arabidopsis *BRASSINAZOLE-RESISTANT1 (BZR1),* one of the main regulators of the brassinosteroid signaling pathway [33]. In Arabidopsis, brassinosteroid signaling inhibits the floral transition and promotes vegetative growth. Furthermore, brassinosteroid-deficient mutants cause a strong delay in days to flowering [64]. Lastly, another QTL was found on Vu03 near *Vigun03g104200,* an ortholog of the soybean *E6* gene, which affects both flowering and maturity [65,66]. *E6* plays an important role in the long-juvenile trait (delayed flowering) in soybean [67].

### Plant productivity traits

Pod load score, dry pod weight and dry fodder weight are important traits related to plant productivity. A cowpea genotype with high pod load (i.e., low pod load score) demonstrates high grain yield potential. High pod load is a result of a high number of pods per plant, which is an indication of low rate of flower abortion. Generally, low flower abortion is associated with resistance to insect attack (typically flower thrips or maruca) and high night temperatures (>18°C) [68]. Negative correlations have been identified between pod load score and dry pod weight, as well as between pod load score and number of pods per plant, and between pod load score and grain yield [69]. The negative correlation between pod load score and grain yield holds true as long as no attack by pod sucking bugs occurs, and in such a situation selection of high grain yielding genotypes can be made using pod load score. However, there are high positive correlations between dry pod weight and grain yield. In some studies, dry pod weight has been negatively correlated with dry fodder weight [70,71] except for some lines with dual purpose characteristics [72,73].

We identified one major QTL associated with pod load score, dry pod weight and dry fodder weight. Seven genes encoding members of the CNGC family proteins were located within the QTL region: four of them (*Vigun04g039300*, *Vigun04g039400*, *Vigun04g039800* and *Vigun04g039900*) are orthologs of the Arabidopsis *CNGC20* gene, also called *CNBT1* (cyclic nucleotide-binding transporter 1), which is involved in the response to nematodes [34,74]. The other three (*Vigun04g039500, Vigun04g039600* and *Vigun04g039700*) are orthologs of the Arabidopsis *CNGC19* gene, which is involved in herbivore response [34,36]. Interestingly, this region corresponds to the major *Rk* locus for root-knot nematode resistance identified in cowpea [75,76]. However, the authors are not aware of any nematode infestation or herbivore damage at the Minjibir field location. Members of the CNGC family of proteins have also been involved in plant tolerance to heavy metals [37]. *CNGC20* and *CNGC19* play essential roles in the responses to biotic and abiotic stresses and thereby may influence plant productivity. Further studies are necessary clarify any possible role of these cowpea homologs in regulating pod load score, dry pod weight and dry fodder weight.

## SUPPLEMENTARY MATERIALS

**Figure S1.** Exploration of the optimal number of subpopulations in the UCR Minicore.

**Figure S2.** Linkage disequilibrium decay for the eleven cowpea chromosomes.

**Figure S3.** Principal component analysis of the UCR Minicore.

**Table S1.** Information on the 368 accessions in the UCR Minicore.

**Table S2.** SNP data from accessions in the UCR Minicore.

**Table S3.** Significant SNPs for DTF in the five different environments.

**Table S4.** Significant SNPs for pod load score, dry pod weight and dry fodder weight.

## FUNDING

This research was funded by the Feed the Future Innovation Lab for Climate Resilient Cowpea (USAID Cooperative Agreement AID-OAA-A-13-00070) and the National Science Foundation BREAD project “Advancing the Cowpea Genome for Food Security” (NSF IOS-1543963). M.C., I.C. and V.C. are supported by National Funds from FCT - Portuguese Foundation for Science and Technology, under the project grant number UIDB/04033/2020.

## ACKNOWLEDGMENTS

The authors acknowledge: Jasmine Dixon for assistance with retrieval of cowpea accessions from storage at UC Riverside; Teresa Cicero and Domenicantonio Vinci for collecting and providing seed of accessions from Vazzano (Italy); Dr. Exequiel Ezcurra for helpful input on the introduction of cowpea into California; Dr. Paul Gepts for helpful discussions; Steve Wanamaker for database assistance; and the University of Southern California Molecular Genomics Core team for iSelect genotyping services.

## CONFLICT OF INTEREST

The authors declare no conflict of interest.

## Notes

### Competing Interest Statement

The authors have declared no competing interest.

## REFERENCES

1. Boukar, O.; Togola, A.; Chamarthi, S.; Belko, N.; Ishikawa, H.; Suzuki, K.; Fatokun, C. Cowpea [Vigna unguiculata (L.) Walp.] Breeding. In Advances in Plant Breeding Strategies: Legumes, Springer: 2019; pp. 201–243.

2. Knox, J.; Hess, T.; Daccache, A.; Wheeler, T. Climate change impacts on crop productivity in Africa and South Asia. Environmental Research Letters 2012, 7, 034032.

3. Müller, C.; Robertson, R.D. Projecting future crop productivity for global economic modeling. Agricultural Economics 2014, 45, 37–50.

4. Maréchal, R.; Mascherpa, J.-M.; Stainier, F. Etude taxonomique d’un groupe complexe d’especes des genres Phaseolus et Vigna (Papilionaceae) sur la base donnees morphologiques et polliniques, traitees par l’analyse informatique. Boissiera 1978, 28, 2–7.

5. Pasquet, R.S. Morphological study of cultivated cowpea *Vigna unguiculata* (L) Walp: importance of ovule number and definition of cv gr *Melanophthalmus*. Agronomie-Sciences des Productions Vegetales et de l’Environnement 1998, 18, 61–70.

6. Xu, P.; Wu, X.; Muñoz-Amatriaín, M.; Wang, B.; Wu, X.; Hu, Y.; Huynh, B.L.; Close, T.J.; Roberts, P.A.; Zhou, W. Genomic regions, cellular components and gene regulatory basis underlying pod length variations in cowpea (*V. unguiculata* L. Walp). Plant Biotechnology Journal 2017, 15, 547–557.

7. Dugje, I.; Omoigui, L.; Ekeleme, F.; Kamara, A.; Ajeigbe, H. Farmers’ guide to cowpea production in West Africa. IITA, Ibadan, Nigeria 2009, 20, 12–14.

8. Brown, A. The core collection at the crossroads. Core collections of plant genetic resources 1995, 3–19.

9. Byrne, P.F.; Volk, G.M.; Gardner, C.; Gore, M.A.; Simon, P.W.; Smith, S. Sustaining the future of plant breeding: The critical role of the USDA-ARS National Plant Germplasm System. Crop Science 2018, 58, 451–468.

10. Huynh, B.-L.; Close, T.J.; Roberts, P.A.; Hu, Z.; Wanamaker, S.; Lucas, M.R.; Chiulele, R.; Cissé, N.; David, A.; Hearne, S., et al. Gene Pools and the Genetic Architecture of Domesticated Cowpea. The Plant Genome 2013, 6, 0, doi:10.3835/plantgenome2013.03.0005.

11. Muchero, W.; Diop, N.N.; Bhat, P.R.; Fenton, R.D.; Wanamaker, S.; Pottorff, M.; Hearne, S.; Cisse, N.; Fatokun, C.; Ehlers, J.D., et al. A consensus genetic map of cowpea *[Vigna unguiculata* (L) Walp.] and synteny based on EST-derived SNPs. Proceedings of the National Academy of Sciences of the United States of America 2009, 106, 18159–18164, doi:10.1073/pnas.0905886106.

12. Xiong, H.; Shi, A.; Mou, B.; Qin, J.; Motes, D.; Lu, W.; Ma, J.; Weng, Y.; Yang, W.; Wu, D. Genetic diversity and population structure of cowpea (*Vigna unguiculata* L. Walp). PloS one 2016, 11, e0160941.

13. Fatokun, C.; Girma, G.; Abberton, M.; Gedil, M.; Unachukwu, N.; Oyatomi, O.; Yusuf, M.; Rabbi, I.; Boukar, O. Genetic diversity and population structure of a mini-core subset from the world cowpea (*Vigna unguiculata* (L.) Walp.) germplasm collection. Scientific Reports 2018, 8, 16035, doi:10.1038/s41598-018-34555-9.

14. Carvalho, M.; Muñoz-Amatriaín, M.; Castro, I.; Lino-Neto, T.; Matos, M.; Egea-Cortines, M.; Rosa, E.; Close, T.; Carnide, V. Genetic diversity and structure of Iberian Peninsula cowpeas compared to world-wide cowpea accessions using high density SNP markers. BMC Genomics 2017, 18, 1–9.

15. Muñoz-Amatriaín, M.; Mirebrahim, H.; Xu, P.; Wanamaker, S.I.; Luo, M.; Alhakami, H.; Alpert, M.; Atokple, I.; Batieno, B.J.; Boukar, O. Genome resources for climate-resilient cowpea, an essential crop for food security. The Plant Journal 2017, 89, 1042–1054.

16. Herniter, I.A.; Muñoz-Amatriaín, M.; Lo, S.; Guo, Y.-N.; Close, T.J. Identification of Candidate Genes Controlling Black Seed Coat and Pod Tip Color in Cowpea (*Vigna unguiculata* [L.] Walp). G3: Genes, Genomes, Genetics 2018, g3. 200521.202018.

17. Herniter, I.A.; Lo, R.; Muñoz-Amatriaín, M.; Lo, S.; Guo, Y.-N.; Huynh, B.-L.; Lucas, M.; Jia, Z.; Roberts, P.A.; Lonardi, S., et al. Seed Coat Pattern QTL and Development in Cowpea (*Vigna unguiculata* [L.] Walp.). Frontiers in Plant Science 2019, 10, doi:10.3389/fpls.2019.01346.

18. Lo, S.; Muñoz-Amatriaín, M.; Hokin, S.A.; Cisse, N.; Roberts, P.A.; Farmer, A.D.; Xu, S.; Close, T.J. A genome-wide association and meta-analysis reveal regions associated with seed size in cowpea *[Vigna unguiculata* (L.) Walp]. Theoretical and Applied Genetics 2019, 132, 3079–3087.

19. Miesho, B.; Hailay, M.; Msiska, U.; Bruno, A.; Malinga, G.M.; Obia Ongom, P.; Edema, R.; Gibson, P.; Rubaihayo, P.; Kyamanywa, S. Identification of candidate genes associated with resistance to bruchid (*Callosobruchus maculatus*) in cowpea. Plant Breeding 2019, 138, 605–613.

20. Steinbrenner, A.D.; Muñoz-Amatriaín, M.; Chaparro, A.F.; Aguilar-Venegas, J.M.; Lo, S.; Okuda, S.; Glauser, G.; Dongiovanni, J.; Shi, D.; Hall, M. A receptor-like protein mediates plant immune responses to herbivore-associated molecular patterns. Proceedings of the National Academy of Sciences 2020, 117, 31510–31518.

21. Lo, S.; Parker, T.; Muñoz-Amatriaín, M.; Berny-Mier y Teran, J.C.; Jernstedt, J.; Close, T.J.; Gepts, P. Genetic, anatomical, and environmental patterns related to pod shattering resistance in domesticated cowpea (*Vigna unguiculata* [L.] Walp). Journal of Experimental Botany under review

22. Muchero, W.; Roberts, P.A.; Diop, N.N.; Drabo, I.; Cisse, N.; Close, T.J.; Muranaka, S.; Boukar, O.; Ehlers, J.D. Genetic architecture of delayed senescence, biomass, and grain yield under drought stress in cowpea. PloS One 2013, 8, e70041, doi:10.1371/journal.pone.0070041.

23. Lonardi, S.; Muñoz-Amatriaín, M.; Liang, Q.; Shu, S.; Wanamaker, S.I.; Lo, S.; Tanskanen, J.; Schulman, A.H.; Zhu, T.; Luo, M.-C., et al. The genome of cowpea (*Vigna unguiculata* [L.] Walp.). The Plant Journal 2019, 98, 767–782, doi:10.1111/tpj.14349.

24. Pritchard, J.K.; Stephens, M.; Donnelly, P. Inference of population structure using multilocus genotype data. Genetics 2000, 155, 945–959.

25. Evanno, G.; Regnaut, S.; Goudet, J. Detecting the number of clusters of individuals using the software STRUCTURE: a simulation study. Molecular Ecology 2005, 14, 2611–2620.

26. Earl, D.A. STRUCTURE HARVESTER: a website and program for visualizing STRUCTURE output and implementing the Evanno method. Conservation Genetics Resources 2012, 4, 359–361.

27. Bradbury, P.J.; Zhang, Z.; Kroon, D.E.; Casstevens, T.M.; Ramdoss, Y.; Buckler, E.S. TASSEL: software for association mapping of complex traits in diverse samples. Bioinformatics 2007, 23, 2633–2635.

28. Zhang, Z.; Ersoz, E.; Lai, C.-Q.; Todhunter, R.J.; Tiwari, H.K.; Gore, M.A.; Bradbury, P.J.; Yu, J.; Arnett, D.K.; Ordovas, J.M. Mixed linear model approach adapted for genome-wide association studies. Nature Genetics 2010, 42, 355.

29. Benjamini, Y.; Hochberg, Y. Controlling the false discovery rate: a practical and powerful approach to multiple testing. Journal of the Royal Statistical Society: series B (Methodological) 1995, 57, 289–300.

30. Huynh, B.L.; Ehlers, J.D.; Huang, B.E.; Muñoz-Amatriaín, M.; Lonardi, S.; Santos, J.R.; Ndeve, A.; Batieno, B.J.; Boukar, O.; Cisse, N. A multi-parent advanced generation inter-cross (MAGIC) population for genetic analysis and improvement of cowpea (*Vigna unguiculata* L. Walp.). The Plant Journal 2018, 93, 1129–1142.

31. Lo, S.; Muñoz-Amatriaín, M.; Boukar, O.; Herniter, I.; Cisse, N.; Guo, Y.-N.; Roberts, P.A.; Xu, S.; Fatokun, C.; Close, T.J. Identification of QTL controlling domestication-related traits in cowpea (*Vigna unguiculata* L. Walp). Scientific Reports 2018, 8, 6261.

32. Marrocco, K.; Zhou, Y.; Bury, E.; Dieterle, M.; Funk, M.; Genschik, P.; Krenz, M.; Stolpe, T.; Kretsch, T. Functional analysis of *EID1*, an F-box protein involved in phytochrome A-dependent light signal transduction. The Plant Journal 2006, 45, 423–438, doi:https://doi.org/10.1111/j.1365-313X.2005.02635.x.

33. He, J.-X.; Gendron, J.M.; Yang, Y.; Li, J.; Wang, Z.-Y. The GSK3-like kinase *BIN2* phosphorylates and destabilizes *BZR1*, a positive regulator of the brassinosteroid signaling pathway in Arabidopsis. Proceedings of the National Academy of Sciences 2002, 99, 10185–10190.

34. Jha, S.K.; Sharma, M.; Pandey, G.K. Role of Cyclic Nucleotide Gated Channels in Stress Management in Plants. Current Genomics 2016, 17, 315–329, doi:10.2174/1389202917666160331202125.

35. Hammes, U.Z.; Schachtman, D.P.; Berg, R.H.; Nielsen, E.; Koch, W.; McIntyre, L.M.; Taylor, C.G. Nematode-induced changes of transporter gene expression in Arabidopsis roots. Molecular plant-microbe interactions 2005, 18, 1247–1257.

36. Meena, M.K.; Prajapati, R.; Krishna, D.; Divakaran, K.; Pandey, Y.; Reichelt, M.; Mathew, M.; Boland, W.; Mithöfer, A.; Vadassery, J. The Ca2+ channel *CNGC19* regulates Arabidopsis defense against Spodoptera herbivory. The Plant Cell 2019, 31, 1539–1562.

37. Moon, J.Y.; Belloeil, C.; Ianna, M.L.; Shin, R. Arabidopsis CNGC family members contribute to heavy metal ion uptake in plants. International Journal of Molecular Sciences 2019, 20, 413.

38. Padulosi, S.; Ng, N. Origin, taxonomy, and morphology of *Vigna unguiculata* (L.) Walp. Advances in cowpea research 1997, 1–12.

39. Steele, W. Cowpeas. In “Evolution of Crop Plants”.(NW Simmonds, ed.). Longman Group. London: 1976.

40. Herniter, I.A.; Muñoz-Amatriaín, M.; Close, T.J. Genetic, textual, and archeological evidence of the historical global spread of cowpea (*Vigna unguiculata* [L.] Walp.). Legume Science 2020, 2, e57.

41. Singh, B.; Ishiyaku, M. Brief communication. Genetics of rough seed coat texture in cowpea. Journal of Heredity 2000, 91, 170–174.

42. Castetter, E.F.; Bell, W.H. Pima and Papago Indian Agriculture. Pima and Papago Indian agriculture. 1942.

43. Fornara, F.; de Montaigu, A.; Coupland, G. SnapShot: control of flowering in Arabidopsis. Cell 2010, 141, 550–550. e552.

44. Craufurd, P.; Qi, A.; Summerfield, R.; Ellis, R.; Roberts, E. Development in cowpea (*Vigna unguiculata).* III. Effects of temperature and photoperiod on time to flowering in photoperiod-sensitive genotypes and screening for photothermal responses. Experimental Agriculture 1996, 32, 29–40.

45. Ehlers, J.; Hall, A. Genotypic classification of cowpea based on responses to heat and photoperiod. Crop Science 1996, 36, 673–679.

46. Kobayashi, Y.; Kaya, H.; Goto, K.; Iwabuchi, M.; Araki, T. A pair of related genes with antagonistic roles in mediating flowering signals. Science 1999, 286, 1960–1962.

47. Yamaguchi, A.; Kobayashi, Y.; Goto, K.; Abe, M.; Araki, T. *TWIN SISTER OF FT* (*TSF*) acts as a floral pathway integrator redundantly with *FT*. Plant and Cell Physiology 2005, 46, 1175–1189, doi:10.1093/pcp/pci151.

48. Díaz, A.; Zikhali, M.; Turner, A.S.; Isaac, P.; Laurie, D.A. Copy number variation affecting the *Photoperiod-B1* and *Vernalization-A1* genes is associated with altered flowering time in wheat (*Triticum aestivum*). PloS One 2012, 7, e33234.

49. Nitcher, R.; Distelfeld, A.; Tan, C.; Yan, L.; Dubcovsky, J. Increased copy number at the *HvFT1* locus is associated with accelerated flowering time in barley. Molecular Genetics and Genomics 2013, 288, 261–275.

50. Dieterle, M.; Zhou, Y.-C.; Schäfer, E.; Funk, M.; Kretsch, T. *EID1*, an F-box protein involved in phytochrome A-specific light signaling. Genes & Development 2001, 15, 939–944.

51. Müller, N.A.; Wijnen, C.L.; Srinivasan, A.; Ryngajllo, M.; Ofner, I.; Lin, T.; Ranjan, A.; West, D.; Maloof, J.N.; Sinha, N.R. Domestication selected for deceleration of the circadian clock in cultivated tomato. Nature Genetics 2016, 48, 89–93.

52. Borner, R.; Kampmann, G.; Chandler, J.; Gleißner, R.; Wisman, E.; Apel, K.; Melzer, S. A MADS domain gene involved in the transition to flowering in Arabidopsis. The Plant Journal 2000, 24, 591–599, doi:https://doi.org/10.1046/j.1365-313x.2000.00906.x.

53. Lee, H.; Suh, S.-S.; Park, E.; Cho, E.; Ahn, J.H.; Kim, S.-G.; Lee, J.S.; Kwon, Y.M.; Lee, I. The *AGAMOUS-LIKE 20* MADS domain protein integrates floral inductive pathways in Arabidopsis. Genes & Development 2000, 14, 2366–2376.

54. Mandel, M.A.; Yanofsky, M.F. The Arabidopsis *AGL8* MADS box gene is expressed in inflorescence meristems and is negatively regulated by *APETALA1*. The Plant Cell 1995, 7, 1763–1771.

55. Hempel, F.D.; Weigel, D.; Mandel, M.A.; Ditta, G.; Zambryski, P.C.; Feldman, L.J.; Yanofsky, M.F. Floral determination and expression of floral regulatory genes in Arabidopsis. Development 1997, 124, 3845–3853.

56. Ferrándiz, C.; Gu, Q.; Martienssen, R.; Yanofsky, M.F. Redundant regulation of meristem identity and plant architecture by *FRUITFULL, APETALA1* and *CAULIFLOWER*. Development 2000, 127, 725–734.

57. Melzer, S.; Lens, F.; Gennen, J.; Vanneste, S.; Rohde, A.; Beeckman, T. Flowering-time genes modulate meristem determinacy and growth form in Arabidopsis thaliana. Nature Genetics 2008, 40, 1489–1492, doi:10.1038/ng.253.

58. Levy, Y.Y.; Mesnage, S.; Mylne, J.S.; Gendall, A.R.; Dean, C. Multiple roles of Arabidopsis *VRN1* in vernalization and flowering time control. Science 2002, 297, 243–246.

59. Putterill, J.; Laurie, R.; Macknight, R. It’s time to flower: the genetic control of flowering time. BioEssays 2004, 26, 363–373, doi:https://doi.org/10.1002/bies.20021.

60. Sung, S.; Amasino, R.M. Remembering winter: toward a molecular understanding of vernalization. Annual Review Plant Biology 2005, 56, 491–508.

61. Trevaskis, B.; Bagnall, D.J.; Ellis, M.H.; Peacock, W.J.; Dennis, E.S. MADS box genes control vernalization-induced flowering in cereals. Proceedings of the National Academy of Sciences 2003, 100, 13099–13104.

62. Yan, L.; Loukoianov, A.; Tranquilli, G.; Helguera, M.; Fahima, T.; Dubcovsky, J. Positional cloning of the wheat vernalization gene *VRN1*. Proceedings of the National Academy of Sciences 2003, 100, 6263–6268.

63. Lü, J.; Suo, H.; Yi, R.; Ma, Q.; Nian, H. *Glyma11g13220,* a homolog of the vernalization pathway gene *VERNALIZATION 1* from soybean *[Glycine max* (L.) Merr.], promotes flowering in Arabidopsis thaliana. BMC Plant Biology 2015, 15, 1–12.

64. Li, Z.; Ou, Y.; Zhang, Z.; Li, J.; He, Y. Brassinosteroid signaling recruits histone 3 lysine-27 demethylation activity to *FLOWERING LOCUS C* chromatin to inhibit the floral transition in Arabidopsis. Molecular Plant 2018, 11, 1135–1146.

65. Li, X.; Fang, C.; Xu, M.; Zhang, F.; Lu, S.; Nan, H.; Su, T.; Li, S.; Zhao, X.; Kong, L., et al. Quantitative trait locus mapping of soybean maturity gene *E6*. Crop Science 2017, 57, 2547–2554, doi:https://doi.org/10.2135/cropsci2017.02.0106.

66. Sedivy, E.J.; Akpertey, A.; Vela, A.; Abadir, S.; Khan, A.; Hanzawa, Y. Identification of non-pleiotropic loci in flowering and maturity control in soybean. Agronomy 2020, 10, 1204.

67. Bonato, E.R.; Vello, N.A. *E6*, a dominant gene conditioning early flowering and maturity in soybeans. Genetics and Molecular Biology 1999, 22, 229–232.

68. Patel, P.; Hall, A. Genotypic variation and classification of cowpea for reproductive responses to high temperature under long photoperiods. Crop Science 1990, 30, 614–621.

69. Garcia-Oliveira, A.L.; Zate, Z.Z.; Olasanmi, B.; Boukar, O.; Gedil, M.; Fatokun, C. Genetic dissection of yield associated traits in a cross between cowpea and yard-long bean (*Vigna unguiculata* (L.) Walp.) based on DArT markers. Journal of Genetics 2020, 99, 1–13.

70. Samireddypalle, A.; Boukar, O.; Grings, E.; Fatokun, C.A.; Kodukula, P.; Devulapalli, R.; Okike, I.; Blümmel, M. Cowpea and groundnut haulms fodder trading and its lessons for multidimensional cowpea improvement for mixed crop livestock systems in West Africa. Frontiers in Plant Science 2017, 8, 30.

71. Singh, B.; Ajeigbe, H.A.; Tarawali, S.A.; Fernandez-Rivera, S.; Abubakar, M. Improving the production and utilization of cowpea as food and fodder. Field Crops Research 2003, 84, 169–177.

72. Boukar, O.; Fatokun, C.A.; Huynh, B.-L.; Roberts, P.A.; Close, T.J. Genomic tools in cowpea breeding programs: status and perspectives. Frontiers in Plant Science 2016, 7, 757.

73. Timko, M.P.; Singh, B. Cowpea, a multifunctional legume. In Genomics of tropical crop plants, Springer: 2008; pp. 227–258.

74. Kaupp, U.B.; Seifert, R. Cyclic nucleotide-gated ion channels. Physiological Reviews 2002, 82, 769–824.

75. Huynh, B.-L.; Matthews, W.C.; Ehlers, J.D.; Lucas, M.R.; Santos, J.R.; Ndeve, A.; Close, T.J.; Roberts, P.A. A major QTL corresponding to the Rk locus for resistance to root-knot nematodes in cowpea (*Vigna unguiculata* L. Walp.). Theoretical and Applied Genetics 2016, 129, 87–95.

76. Ndeve, A.D.; Santos, J.R.; Matthews, W.C.; Huynh, B.L.; Guo, Y.-N.; Lo, S.; Muñoz-Amatriaín, M.; Roberts, P.A. A novel root-knot nematode resistance QTL on chromosome Vu01 in Cowpea. G3: Genes, Genomes, Genetics 2019, 9, 1199–1209.

