## Supplemental Figure S1 for "The UCR Minicore: a valuable resource for cowpea research and breeding"

**Figure S1. Exploration of the optimal number of subpopulations (*K*).** (A) Estimated log probability of the data for each *K* between 1 and 10 (top plot). The same data was plotted for *K* values between 1 and 9 (bottom) as the large standard error for *K*=10 obscures the trend in the data. (B) Δ*K* values as a function of *K*. Plots A (top) and B were generated with Structure Harvester (Earl et al. 2012).

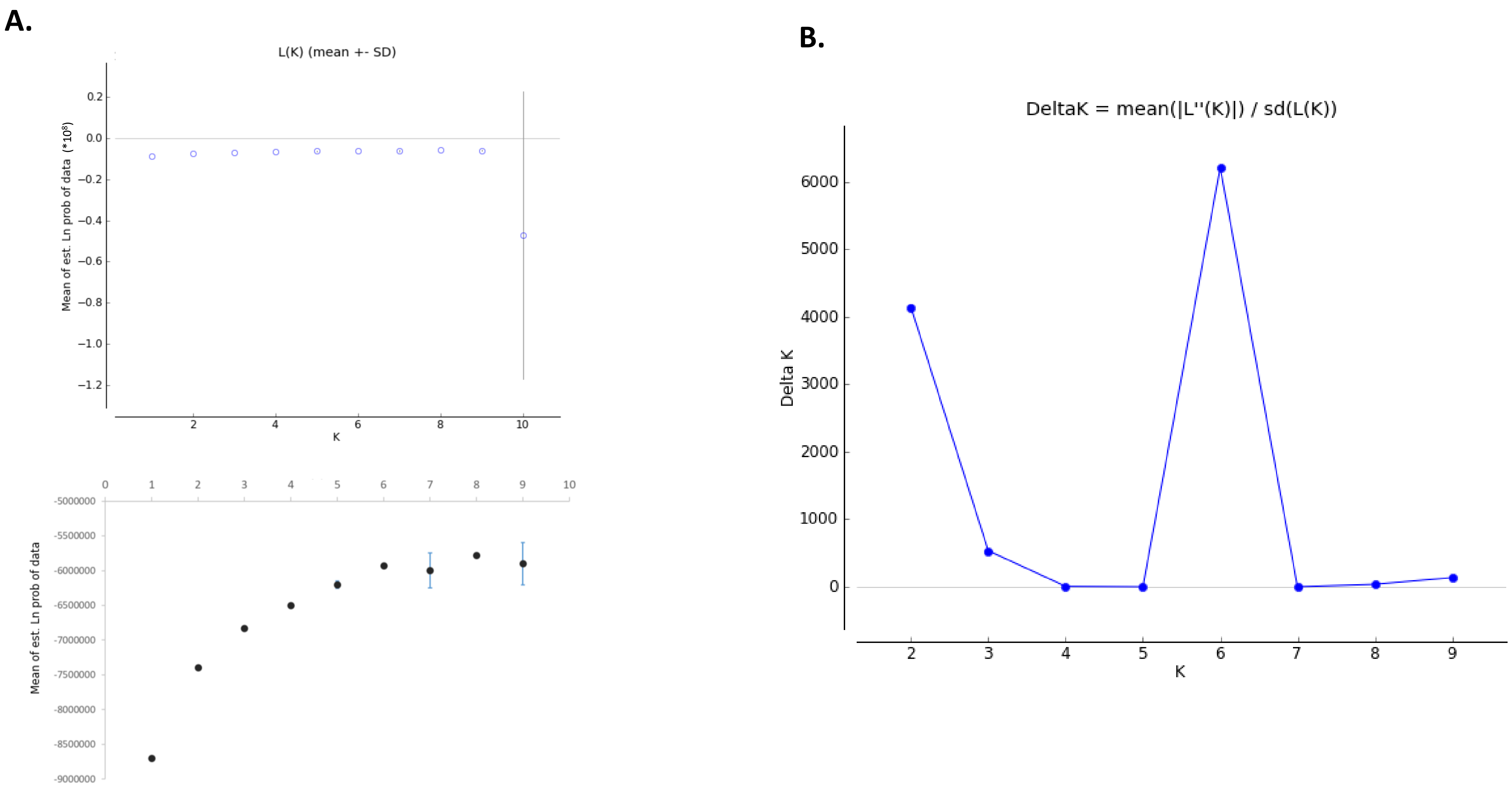
