## Supplemental Figure S2 for "The UCR Minicore: a valuable resource for cowpea research and breeding"

**Figure S2. Linkage disequilibrium decay for the eleven cowpea chromosomes.** The decay of $r^2$ over physical distance (in kb) is shown for Vu01 to Vu11. The black dots indicate the $r^2$ values of all SNP pairs within each chromosome.  The significant threshold is represented as a red horizontal  line in each chromosome and the $r^2$ values falling to the critical threshold are indicated as green vertical lines.

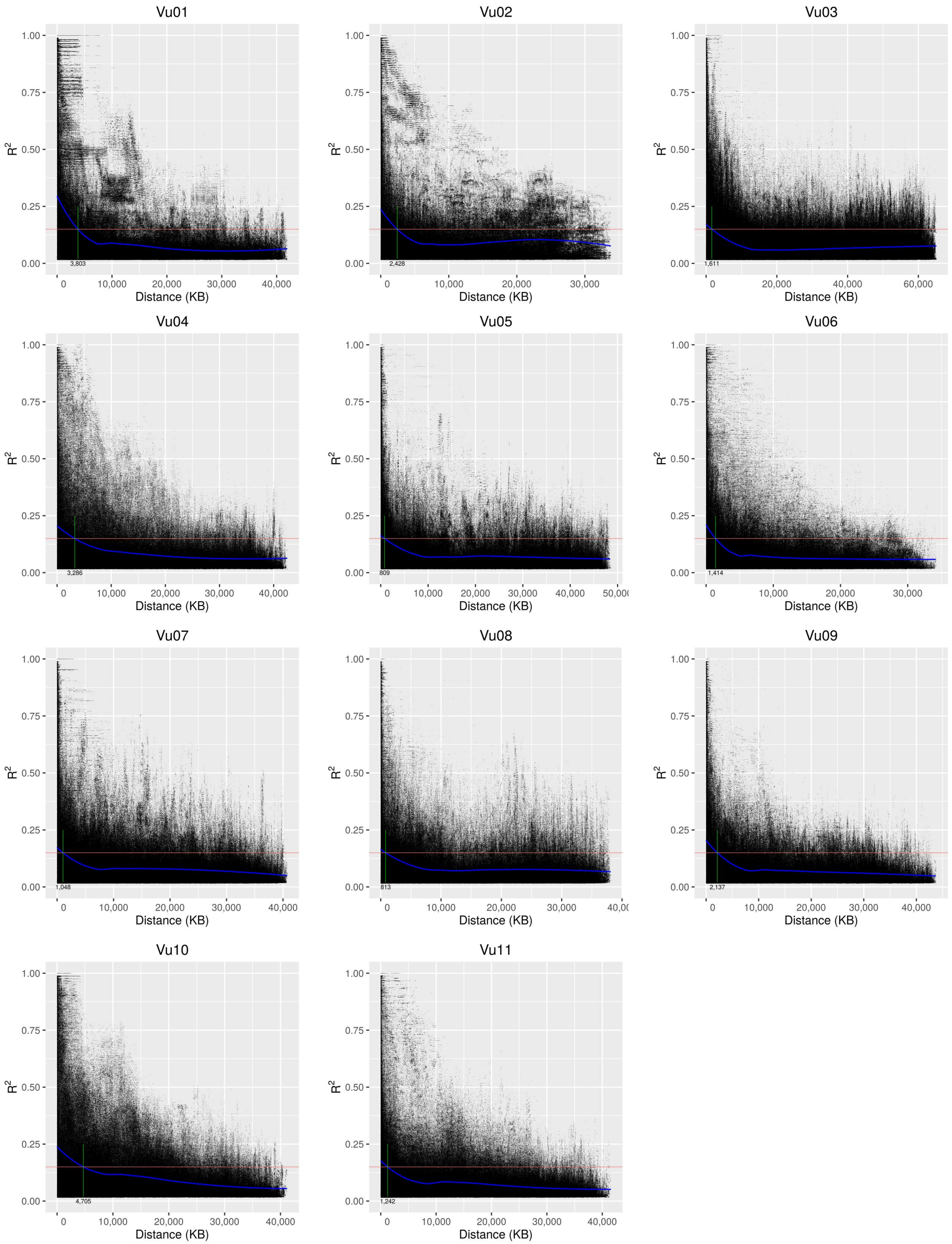
