## Supplemental Figure S3 for "The UCR Minicore: a valuable resource for cowpea research and breeding"

**Figure S3. Principal component analysis of the UCR Minicore.** The first three principal components are shown, with accessions colored by the result of STRUCTURE for *K*=6. Accessions with a membership coefficients < 0.8 were considered “admixed”.

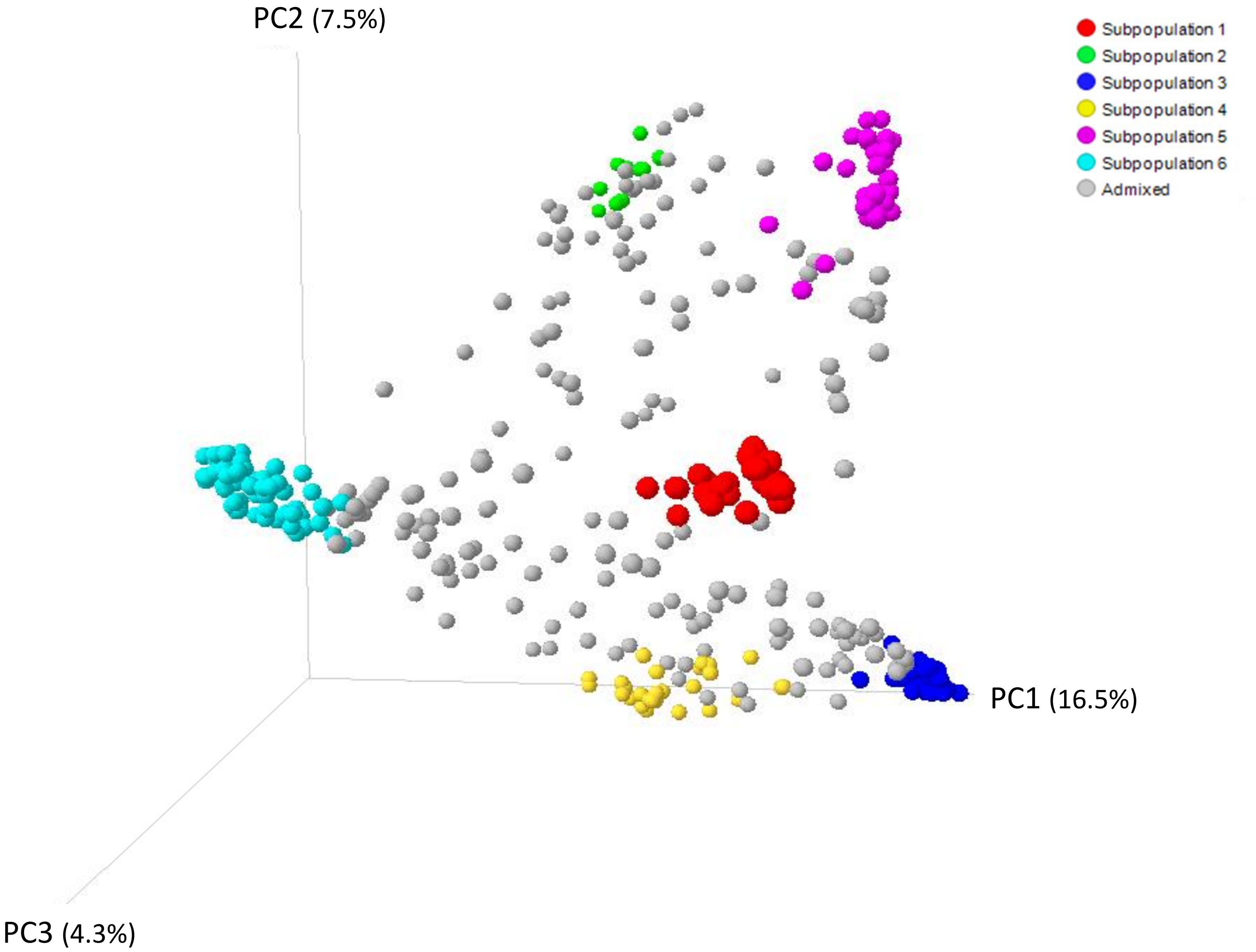
